# The persistent homology of mitochondrial ATP synthases

**DOI:** 10.1101/2022.09.13.506888

**Authors:** Savar D. Sinha, Jeremy G. Wideman

## Abstract

While mitochondrial ATP synthase has been thoroughly studied in animals and fungi, relatively little is known about the structures of protists. Among those that have been studied, protist ATP synthases possess divergent structures distinct from those of yeast or animals. Therefore, we aimed to clarify the subunit composition and evolution of ATP synthase across all major eukaryotic lineages. We used sensitive homology detection methods and molecular modelling tools to demonstrate the persistence of a near-complete ancestral set of 17 subunits in most major eukaryotic taxa even despite major divergence. These data demonstrate that most eukaryotes possess an ancestral-like ATP synthase structure similar to those of animals, fungi, and plants, but a number have diverged drastically (e.g., ciliates, myzozoans, euglenozoans, and likely retarians and heteroloboseans). In addition, we identified the first synapomorphy of the SAR (stramenopile, alveolate, rhizaria) supergroup – a ~1 billion-year-old gene fusion between ATP synthase stator subunits. Our comparative approach highlights the persistence of ancestral subunits even amidst major structural changes. We conclude by urging that more ATP synthase structures (e.g., from jakobids, heteroloboseans, stramenopiles, rhizarians) are needed to provide a complete picture of the evolution of structural diversity of this ancient and essential complex.

## Introduction

Mitochondria arose from an alphaproteobacterial endosymbiont over 1.5 billion years ago (1–3). Despite its prokaryotic origins, only ~10-20% of the mitochondrial proteome is directly descended from homologs in alphaproteobacteria (4). All other mitochondrial proteins were either acquired through horizontal gene transfer (HGT) from other bacteria or originated *de novo* in the eukaryotic nuclear genome during eukaryogenesis (5). The most famous and perhaps the most important function of mitochondria is their involvement in the formation of a proton gradient via Complexes I-IV of the electron transport chain (ETC) and the ultimate production of ATP via ATP synthase (Complex V) (6). The eukaryotic versions of the ETC and ATP synthase retain many of the basic features of their prokaryotic counterparts; however, eukaryotes have accreted numerous supernumerary protein subunits (7). It is currently unclear which of these additional components are ancestral to eukaryotes and which are lineage specific.

All studied mitochondrial ATP synthases form dimers, whereas their counterparts in bacteria and chloroplasts do not (8). It has been proposed that the eukaryotic supernumerary ATP synthase subunits evolved to facilitate their dimerization as well as the formation and maintenance of cristae (9, 10). In fact, variation in ATP synthase dimerization angle correlates well with variations in cristae morphology and ultrastructure, providing further support linking molecular structure to cell biological variation (11). ATP synthases have been categorized into four different types. Animals and fungi possess the canonical Type I ATP synthase and lamellar cristae (12, 13). Chlorophycean algae (e.g., *Chlamydomonas* and *Polytomella*) contain Type II ATP synthases and club-shaped cristae (9). Two groups of related protists (ciliates and apicomplexans) contain Type III ATP synthases. The ciliate *Tetrahymena thermophila* has helical tubular cristae, whereas the apicomplexan *Toxoplasma gondii*, has a bulbous cristae morphology (14, 15). Euglenozoan protists like *Trypanosoma* and *Euglena* contain Type IV ATP synthases and discoid cristae (16–18).

Early investigations of Type I ATP synthases demonstrated that the same set of 17 subunits compose the dimeric complexes of both animals and fungi (7, 19) (Figure 1). Type I ATP synthases comprise two sectors which can be separated biochemically. The soluble F_1_ sector is composed of five subunits (α, β, γ, δ, and ε) and is the site of ATP synthesis from ADP and P_i_ via rotation of the γ-subunit. The membrane bound F_o_ sector comprises the peripheral stalk and proton half-channels. The a-subunit facilitates the transfer of protons down their concentration gradient via the rotation of the c-ring, an oligomer of 10 c-subunits in most eukaryotes (20). The remaining subunits (b, d, e, f, g, h, i/j, k, 8, and OSCP) compose the peripheral stalk, which links the F_o_ to the F_1_ sector. Subunits f, e, g, i/j and k are putatively involved in dimerization (12). Many of these subunits are present in Types II, III, and IV ATP synthases; however, the ancestry of several subunits in these complexes remains unresolved. Furthermore, due to the extremely short length (<100 aa) of many of these protein subunits, an extensive comparative genomics survey including representatives from across eukaryotic diversity has not been undertaken.

**Figure 1.**
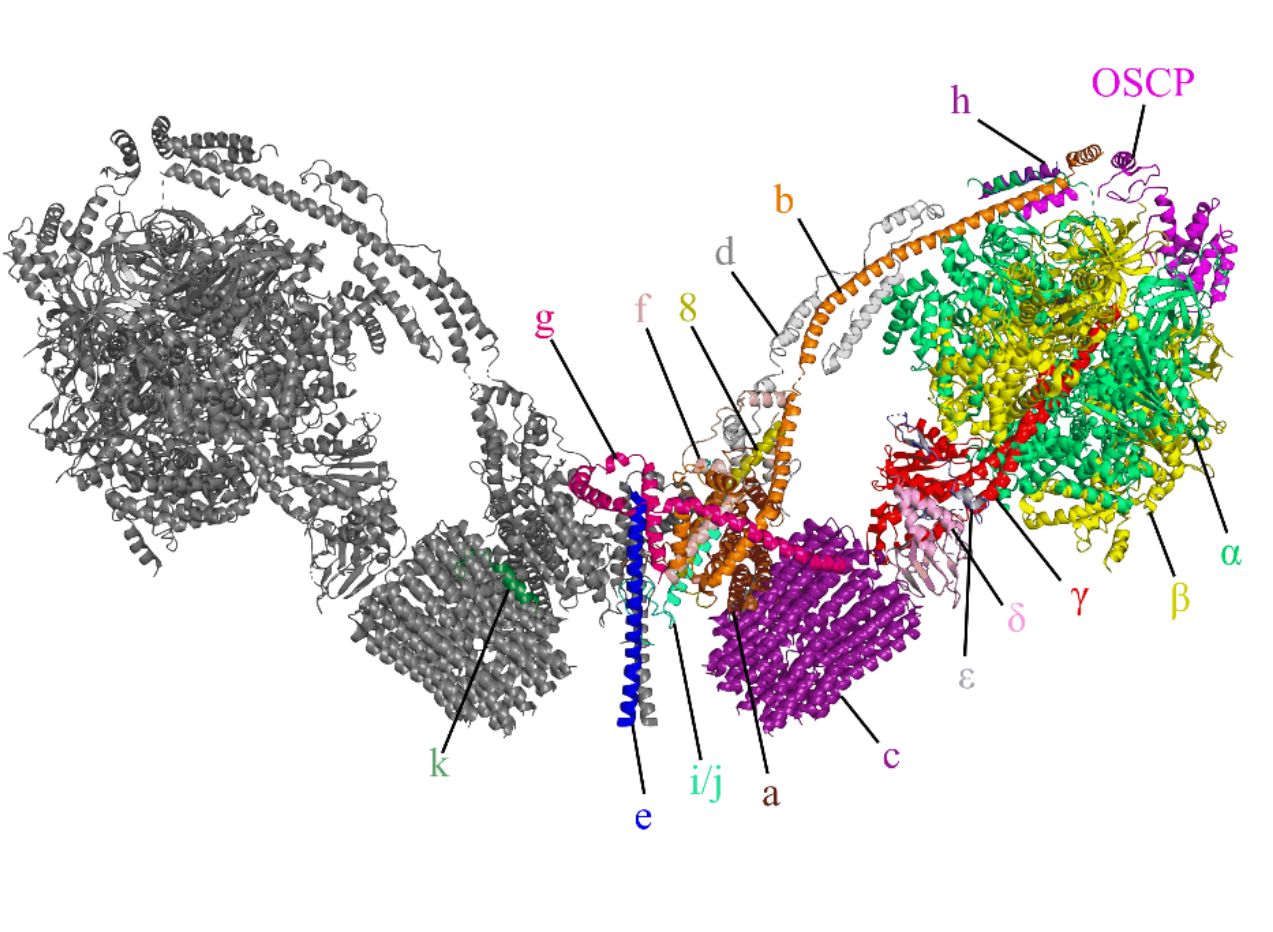
Structure of *Saccharomyces cerevisiae* Type I F_1_F_o_ ATP synthase dimer. The Cryo-EM Type I structure from yeast represents the typical ATP synthase structure found across eukaryotes (PDB: 6B8H). The F_1_ sector comprises the catalytic head of the complex including the αβ hexamer and the central rotor γ + δ, and ε. The F_o_ sector comprises the peripheral stalk (8, b, d, f, h, OSCP), H^+^ channel (a and c), and intermembrane dimerization subunits (e, g, i/j, k).

Here, we use diverse bioinformatic tools to identify ATP synthase subunits in a broad diversity of 219 eukaryotic predicted proteomes. We identify several divergent proteins in Type II, III, and IV ATP synthases as divergent homologues of canonical Type I subunits and conclude that the ATP synthase present in the Last Eukaryotic Common Ancestor (LECA) contained the 17 canonical Type I ATP synthase subunits. Furthermore, we show that most ATP synthase subunits are conserved across most eukaryotic lineages and suggest that novel subunits tend to be accreted and do not usually replace ancestral subunits. Finally, we identified a ~1 billion-year-old gene fusion between subunits b and h of the peripheral stalk in the SAR supergroup, the very first synapomorphy of this lineage. Collectively, these data provide a broad picture of the evolution of eukaryotic ATP synthases.

## Results and Discussion

### The ancestral eukaryotic ATP synthase contained all 17 subunits shared by animal and fungal complexes

To determine the ancestral subunit complement of eukaryotic ATP synthase, we first collected representative protein sequences from eukaryotic species with solved or previously characterized structures (Dataset S1). We used these sequences as BLAST (21, 22) queries into an expanded version of The Comparative Set (TCS), which is a subset of EukProt v3 (23) predicted proteomes selected based on their phylogenetic importance and BUSCO completion (See Methods for more details and Dataset S1 for a complete list of proteomes used in this study). We added a few extra species not included in the TCS for a total of 219 predicted proteomes. Top hits with E-values below a threshold of 0.01 were used as reciprocal BLAST queries (21, 22) into predicted proteomes of species with solved ATP synthase structures. Orthologues were considered validated if *bona fide* ATP synthase subunits were retrieved as top hits of reciprocal searches. Orthologue sets of each ATP synthase subunit were then aligned to build Hidden Markov Models (HMMs) to use in HMMER (24, 25) searches to identify more divergent orthologues. In several extreme cases, divergent orthologues could not be validated using reciprocal searching, and structural comparisons using Phyre2 (26), SWISS-MODEL (27) and AlphaFold2 (28) were used to infer orthology (See Figures S1-S7). Using a mixture of these approaches, we identified all 17 canonical animal/fungal ATP synthase subunits across diverse lineages spanning the entire tree of eukaryotes (Figure 2). From these data we conclude that, under nearly any eukaryotic rooting hypothesis (29–31), the ancestral ATP synthase contained all 17 animal/fungal subunits. Similarly, a complex complement of ATP synthase assembly factors is also retained in most eukaryotic lineages (Figure S8). Certain factors like Factor B and Atp25 were confirmed to be animal- or fungal-specific, respectively. All other subunits are retained among most lineages with notable absences of Atp10 in Chlorophyceans; Atp10, Nca2, TMEM70, and TMEM242 in select alveolate lineages; and TMEM70 across most discicristates.

**Figure 2.**
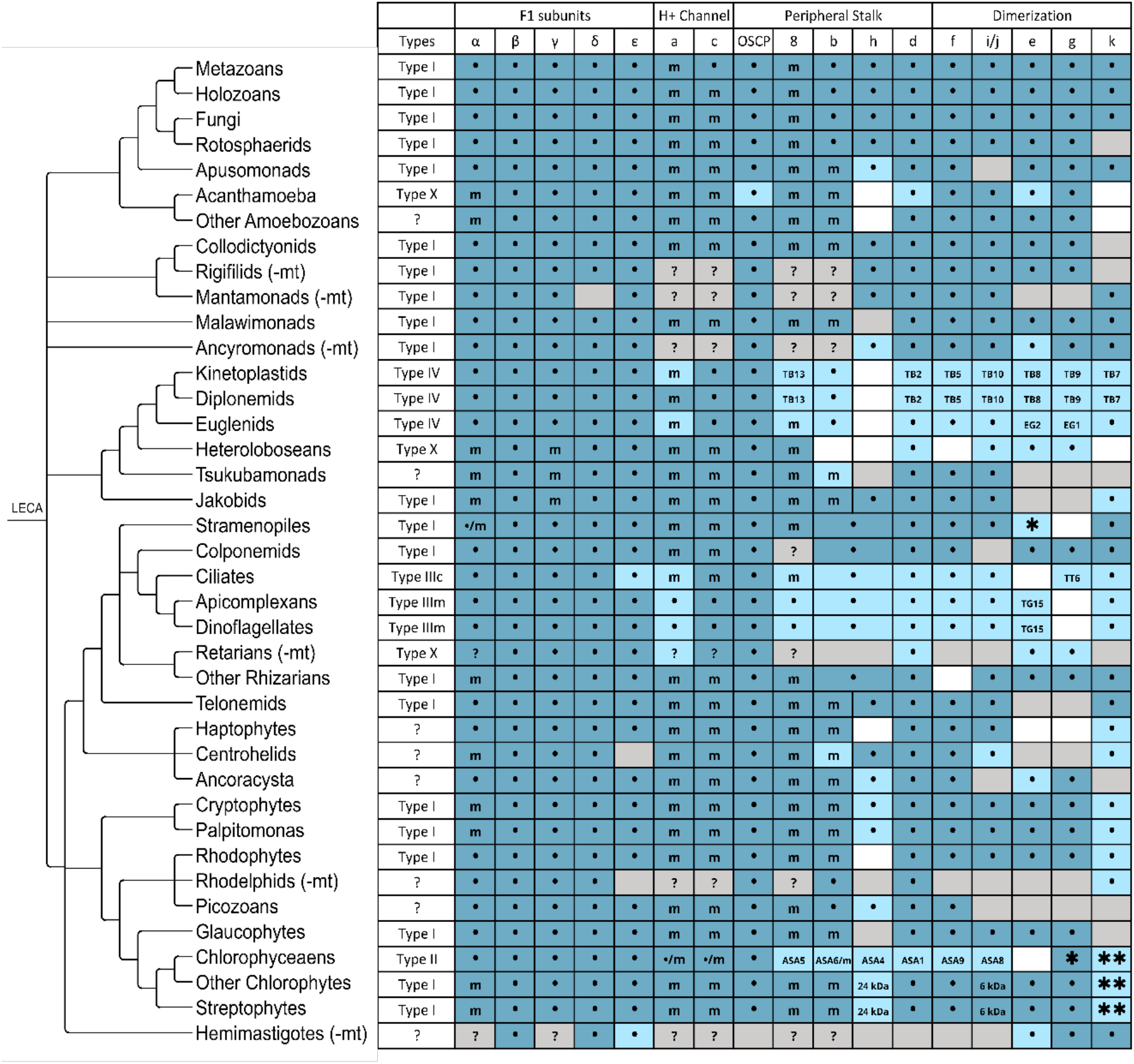
Most eukaryotic ATP synthases retain the core set of 17 ancestral subunits. ATP synthase orthologues were identified in 219 EukProt3 proteomes from major eukaryotic taxa. Subunits were collected via BLAST and HMMER. Dark blue indicates verification via reciprocal BLAST or phmmer (21, 22, 25) and light blue via HHpred or structural modelling (26–28, 64). Grey indicates lack of genomic data. Subunits misidentified as novel are indicated by their species-specific names (e.g. TB5). •/^m^Subunits are nuclear/mitochondrially encoded respectively. *Presence is restricted to certain lineages. **Subunit may no longer associate with ATP synthase. ^(−mt)^Taxa lack published mitochondrial genomes. ^?^Presence in mitochondrial genome is unknown.

### Divergent ATP synthases retain most ancestral protein subunits

While all investigated eukaryotic ATP synthases exhibit dimeric structures (32), one of the most conspicuous structural differences between lineages is their dimer angles (9, 12–16, 18). These dimer angle variations likely influence mitochondrial cristae shape as they correlate with differences in mitochondrial ultrastructure (9). Alongside divergences in cristae morphology and dimer angles, ancestral ATP synthase subunits have diverged so greatly in certain lineages that several subunits were originally mis-identified as novel lineage-specific accretions or replacements (Dataset S2 and (14–17)). Here, we were able to bioinformatically identify several putatively lineage-specific proteins in Type II, III, and IV ATP synthases as orthologues of Type I subunits (Figure 2).

Unlike other green algae, Chlorophycean Type II ATP synthases like that of *Polytomella parva* have divergent structures (33). Type II ATP synthases possess an enlarged peripheral stalk with a smaller dimer angle of 56° (compared to 86° in yeast), resulting in club-shaped cristae (9). Type II ATP synthases include several F_o_ subunits widely conserved across Chlorophyceans (ASA1-ASA10) that were originally thought to be lineage-specific, though we identified most as orthologues of ancestral subunits (33, 34). We corroborated previous studies which suggested ASA4 as homologous to subunit h (35). Through HHpred searches and structural comparisons of helix location in assembled complexes, we identified ASA1 as subunit d (Figure S3), ASA5 as subunit 8 (Figure S4), ASA6 as subunit b (Figure S2), ASA8 as subunit i/j (Figure S6), and ASA9 as subunit f (Figure S5). We note that ASA10 matches the location and runs in the opposite direction of a N-terminal helix lost in Chlorophycean subunit a, suggesting a possible partial replacement (Figure S9). HMMER also identified divergent subunit k orthologues in green plants and algae that appear to be absent from the solved Type II complex (Figure 2).

It should be noted however that the composition of Type II ATP synthase is not uniform across chlorophyceans. While all ATP synthase subunits are nuclear encoded in chlamydomonad algae such as *Polytomella parva* and *Chlamydomonas reinhardtii*, our study revealed the presence of subunits a (*atp6*), c (*atp9*), and ASA6 in the mitochondrial genome of *Tetradesmus obliquus*, supporting our identification of ASA6 as the orthologue of subunit b (*atp4*), which is mitochondrially encoded across most taxa. Since no protein comparable to ASA5 or *atp8* was identified in the nuclear genome of *Tetradesmus obliquus*, it is possible that subunit 8 is also mitochondrially encoded. Furthermore, in *Tetradesmus obliquus*, we identified an ortholog of subunit g, which appears to have been lost in most chlamydomonad algae. These observations support the gradual evolution of Type II ATP synthase from a Type I ancestor.

Although the ATP synthases from the alveolate lineages of myzozoans (e.g., *Toxoplasma gondii*) and ciliates (e.g., *Tetrahymena thermophila*) are very divergent from one another, they have been collectively categorized as Type III complexes. To distinguish between the entirely distinct ATP synthases of ciliates and myzozoans (i.e., the clade comprising apicomplexans and dinoflagellates), we have designated them as Type IIIc (ciliate) and Type IIIm (myzozoan) ATP synthases, respectively. In the Type IIIc and Type IIIm complexes, the ATP synthase monomers are both positioned approximately parallel to each other (~0° dimer angles), producing vastly different cristae structures from both one another and other ATP synthases. The tetramers of Type IIIc produce tubular cristae, while the hexamers of Type IIIm form bulbous cristae (14, 15). Both the Type III ATP synthases present a wide range of conserved ancestral subunits; however, Type IIIc contains 12 novel subunits universally conserved across ciliates, while Type IIIm comprises 16 unrelated lineage-specific subunits conserved in all apicomplexans, 12 of which are also conserved in dinoflagellates (Table S1) (14, 15, 36, 37). Through HHpred sequence comparison with alignments of subunits e and g obtained from the 219 EukProt3 genomes and transcriptomes, we identified ATPTG15 as a putative orthologue of subunit e and ATPTT6 as a putative subunit g sequence (Figure 2); however, we were unable to identify a candidate subunit g in apicomplexans or a subunit e in ciliates (see Dataset S1). The widespread conservation of Type III ATP synthases among ciliates and myzozoans indicates that these structural changes occurred early in the evolution of these lineages and have remained relatively stable for over 500 million years (38).

Extensive research on *Trypanosoma brucei* and *Euglena gracilis* has revealed the incredibly divergent structure of euglenozoan Type IV ATP synthases. Similar to the other divergent complexes, the euglenid and kinetoplastid structures possess smaller dimer angles of 45° and 60° respectively, with novel subunits accumulated around the periphery of both the F_1_ and F_o_ sectors (16–18). All three aerobic euglenozoan lineages (kinetoplastids, diplonemids, and euglenids) share 8 lineage-specific ATP synthase subunits, with an additional four present only in euglenids and one specific to kinetoplastids and diplonemids (Table S1). During manuscript preparation, a subset of our bioinformatic predictions were validated by structural and biochemical analyses of the *Trypanosoma brucei* ATP synthase (16, 17). Using HMMsearch, HHpred, and structural validation, we were able verify the putative identities of ancestral subunits originally classified as novel, including ATPTB2 as subunit d (Figure S3), ATPTB5 as subunit f (Figure S5), ATPTB7 as subunit k (Figure S7), ATPTB8/ATPEG2 as subunit e, ATPTB9/ATPEG1 as subunit g (Figure 4), ATPTB10 as subunit i/j (Figure S6), and ATPTB13 as subunit 8 (Figure S4) (16, 17, 39). However, even though we were able to identify many subunits, kinetoplastid subunit b could not be found bioinformatically. Two recent reports identified two unrelated proteins as candidates for kinetoplastid subunit b (16, 40). Our homology searches revealed that both candidate subunit b proteins are universally conserved in sequenced kinetoplastids and diplonemids; however, they bear little resemblance to each other, their euglenid counterpart, or any other subunit b orthologue (Figure 3).

**Figure 3.**
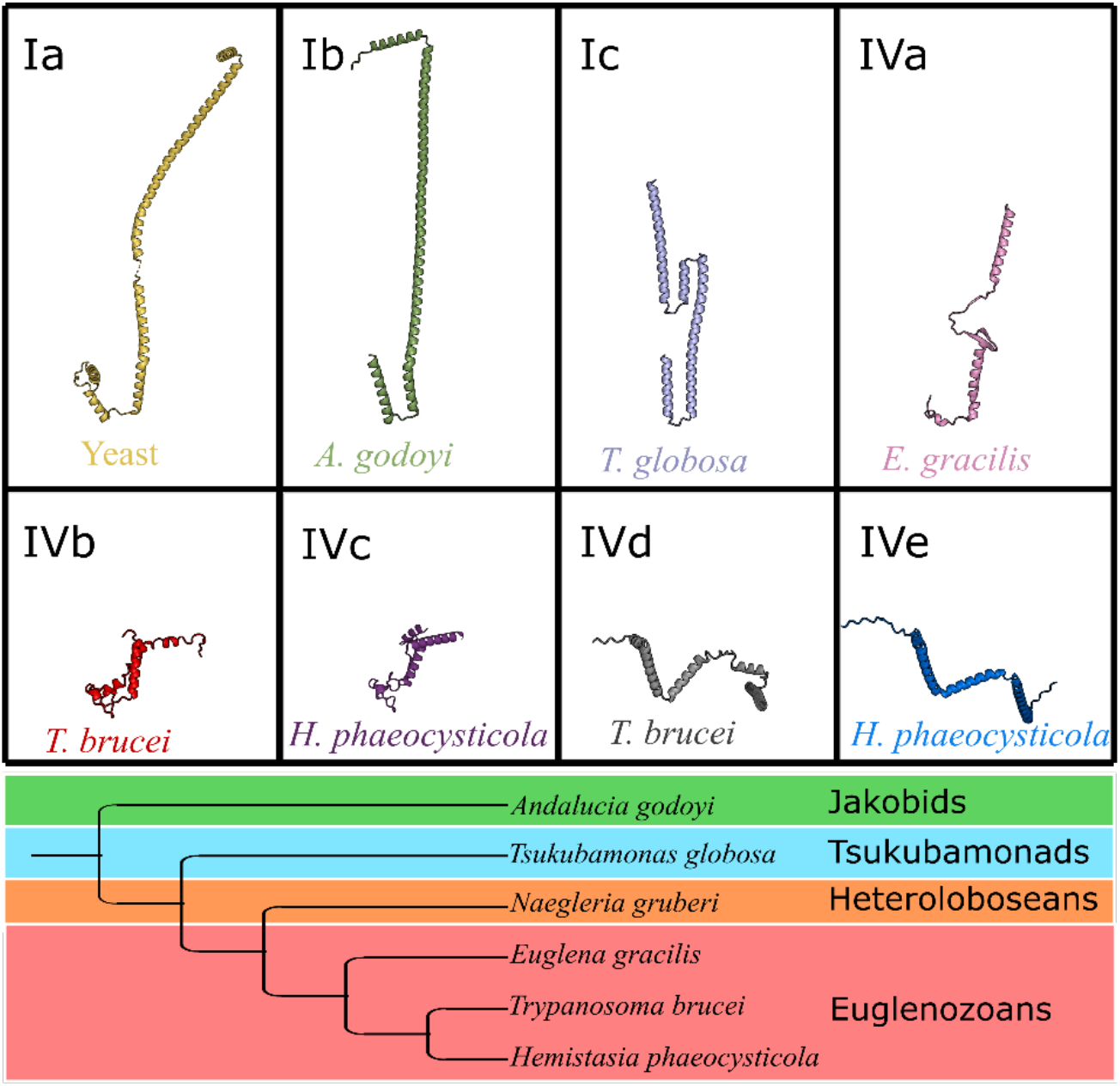
Truncation of subunit b in the transition from Type I to Type IV ATP synthases in the lineage leading to Trypanosomes. The ancestral b-subunit was similar to the b subunit from bacteria and is still similar in Type I ATP synthases. ^Ia^Structure of *Bos taurus* (bovine) subunit b (PDB: 6ZPO). ^Ib^*Andalucia godoyi* subunit b and ^It^*Tsukubamonas globosa* subunit b modelled using AlphaFold2 (28). ^IVa^*Euglena gracilis* subunit b (PDB: 6TDU). Sequences for *Trypanosoma brucei* were obtained from the recent Cryo-EM study (16) and biochemical study (40) used as BLAST searches to identify orthologues from diplonemid *Hemistasia phaeocysticola*. ^IVb^*T. brucei* subunit b modelled using AlphaFold2 (28). ^IVc^ *H. phaeocysticola* ‘Cryo-EM’ subunit b. ^IVd^*T. brucei* subunit b identified biochemically. ^IVe^*H. phaeocysticola* ‘biochemical’ subunit b. Tree shown below depicts the evolution of discobids (23).

**Figure 4.**
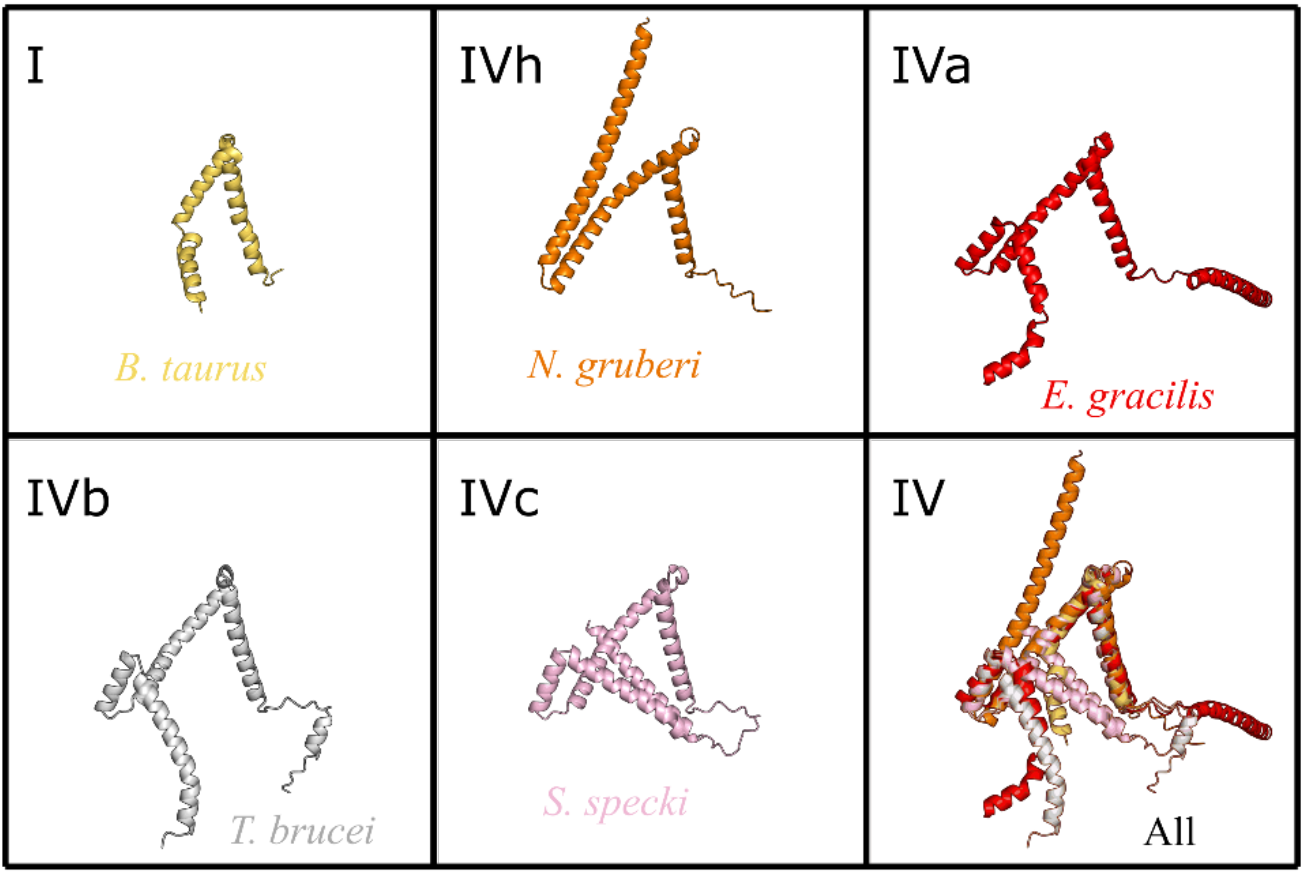
Heterolobosean subunit g confirms the orthology of euglenozoan subunits. ^I^Structure of bovine subunit g. ^IVh^*Naegleria gruberi* subunit g. ^IVa^*Euglena gracilis* subunit g. ^IVb^*Trypanosoma brucei* subunit g. ^IVc^*Sulcionema specki* subunit g. ^IV^Euglenozoan g-subunits overlayed. Structures from *Bos taurus* (PDB: 6ZPO) and *Euglena gracilis* (PDB: 6TDU) were obtained from PDB. Sequence for *Naegleria gruberi* was obtained via HMMER and structurally modelled using AlphaFold2 (25, 28). Structures were aligned through PyMOL2 (65–67).

Other discobids closely related to euglenozoans lack the shared characteristics of the euglenozoan ATP synthase. For example, jakobids, which contain the most gene-rich and bacteria-like mitochondrial genome appear to have more ancestral-like ATP synthase structures and virtually contain a Type I ATP synthase lacking detectable e and g subunits ((41) and Figure 2). Whereas tsukubamonads and heteroloboseans, which branch nearest euglenozoans, may possess ATP synthases of intermediate divergence. While subunits a, c, d, i/j, and 8 are readily identifiable in tsukubomonads and heteroloboseans, other components including subunits h and k were more difficult to identify conclusively. Furthermore, subunit f and a divergent mitochondrially encoded subunit b were identified in *Tsukubamonas globosa*; however, no such subunit was identified in heteroloboseans, suggesting that only structural investigations will identify these universally conserved subunits. Moreover, a euglenozoan-like g-subunit orthologue which structurally resembles ATPEG1 according to SWISS-MODEL, was identified in heteroloboseans (Figure 4), and could provide clues into the formation of the discoid cristae shapes conserved in both heteroloboseans and euglenozoans (17, 18, 42). Finally, the divergence of subunit b in tsukubamonads paired with the inability to identify a heterolobosean subunit b suggests that a gradual divergence of Type IV ATP synthase may have occurred in the discicristate lineage. What the explanation for this divergence might be is still a mystery. Both adaptive and constructive neutral evolutionary scenarios could be at play (43). More intermediate lineages that branch at the base of euglenozoans (e.g., EU17/18 (44)) are required to better understand the evolution of Type IV ATP synthases.

### Absence of evidence is not evidence of absence: small subunits e, f, g, h, and k are likely present but not detected in most predicted proteomes

The recent euglenozoan structural data and ambiguous orthology of subunits e and g in alveolates indicate the requirement of structural data to identify core ATP synthase subunits in divergent structures. While our investigation has uncovered the broad conservation of all ATP synthase subunits in several major eukaryotic lineages, even the most sensitive homology detection tools cannot unambiguously identify orthologues of small divergent subunits (e.g., subunits e, f, g, h, k). Furthermore, since the default parameters of many gene prediction programs eliminate proteins shorter than 100 aa, short ATP synthase subunits are often left unpredicted. Thus, many of these smaller subunits could only be detected in the transcriptomes or raw reads in the SRA database (See Table S2 cells in orange). Since HMM searches could not be conducted on these databases due to computational limitations, some more divergent small subunits may remain unidentified.

In animals and fungi, both subunits e and g are necessary for dimer stability (45). However, we were unable to identify both subunits in numerous lineages. In stramenopiles and haptophytes, we identified putative e subunits among a select few taxa but could not identify dimerization subunits in most species in these lineages. Furthermore, In Type III ATP synthases, no probable subunit g was found in myzozoans and no subunit e was identified in ciliates. Neither subunit e nor g could be identified in jakobids, and in tskubamonads, we were unable to uncover any orthologues of subunit e (Figure 2). Thus, only structural investigations will uncover the dimerization mechanisms of these lineages.

Subunit f is a larger subunit present among almost all taxa investigated, with notable absences in heteroloboseans and rhizarians. Given the presence of divergent subunit f sequences in euglenozoans, it is likely that subunit f is similarly divergent in heteroloboseans. The absence of subunit f in all rhizarians may also indicate minor alterations to the general structure of the rhizarian complex (Figure 2).

### SAR lineage synapomorphy revealed in billion-year-old gene fusion between subunits b and h

The SAR supergroup links stramenopiles, alveolates, and rhizarians together into a single clade identified only by multigene phylogenetics (46). Despite this phylogenetic support, to our knowledge, no shared organismal or molecular traits that characterize the SAR supergroup have been identified. Here, we present the first synapomorphy of the SAR supergroup – a gene fusion of the ATP synthase b and h subunits (Figure 5).

**Figure 5.**
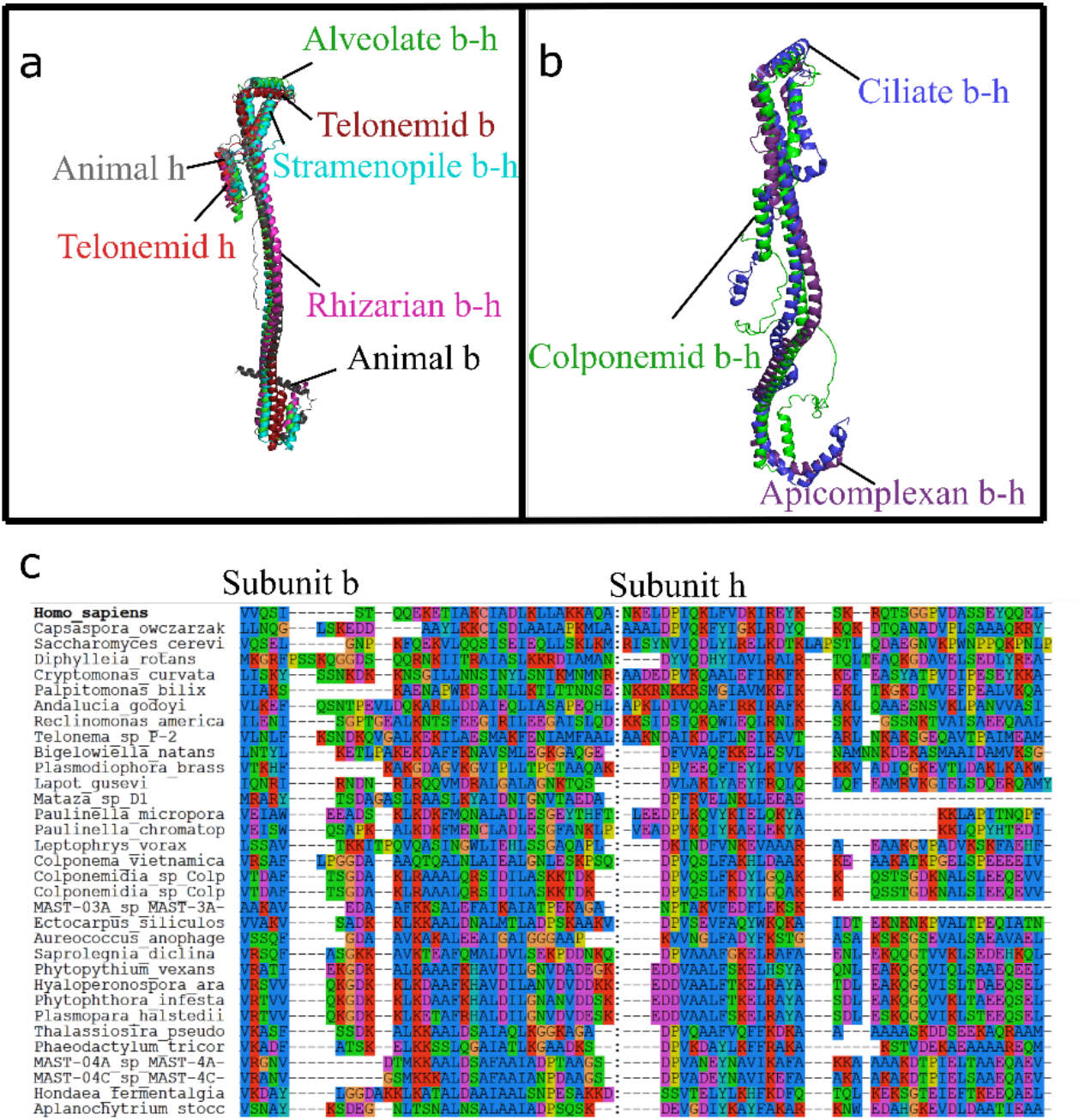
Billion-year-old subunit b-h fusion is a synapomorphy of the eukaryotic clade comprising stramenopiles, alveolates, and rhizarians (SAR). **(a)**. Structural alignment of predicted stramenopile, alveolate, and rhizarian b-h fusions with b and h subunits from Type I structures. Since some SAR ATP synthases are extremely divergent (e.g., Type III from ciliates and myzozoans), we chose minimally divergent representatives from each SAR clade for structural modelling using AlphaFold2 and subsequent alignment via PyMOL2 (28, 65–67). Stramenopile: *Ectocarpus siliculosus* (blue). Alveolate: *Colponema vietnamica* (green). Rhizarian: *Plasmodiophora brassicae* (pink). Telonemid: *Telonema sp. P-2* (red). **(b)**. Type III alveolate ATP synthases retain highly divergent b-h fusion proteins. *C. vietnamica* b-h subunit was structurally aligned to known structures of b subunits from *Tetrahymena thermophila* and *Toxoplasma gondii* (14, 15). **(c)**. SAR b-h subunit align to concatenated b and h subunits from across eukaryotes. N-terminal subunit h mitochondrial targeting sequences were identified using TargetP 2.0 (68) and removed. These trimmed h subunits were appended to b subunits from the same species. These artificially fused b and h subunit sequences were then aligned to b-h subunits from stramenopiles, alveolates, and rhizarians using MUSCLE (63) and visualized using Seaview Version 5 (69). The approximate b-h boundary is represented by a line of colons.

The ancestral b subunit was encoded in the mitochondrial genome of LECA and is still encoded in the mitochondrial genome of many extant eukaryotes, including the sister group to SAR, the telonemids (47). All SAR representatives encode subunit b in their nuclear genome (Figure 2). This means that ~1 billion years ago, prior to the diversification of SAR, subunit b was likely transferred from the mitochondrial to the nuclear genome. In our investigations of SAR b subunits, we noticed consistent C-terminal extensions. HHpred searches and structural modelling revealed the C-terminal portion of SAR b subunits to share homology with h-subunits, which otherwise cannot be identified in any SAR species. Thus, we suggest that upon its transfer to the nucleus, the subunit b gene fused with subunit h, producing the b-h subunit of stramenopiles, alveolates, and rhizarians. These data help explain the h-like structure observed at the C-terminus of Type IIIm subunit b (15).

Given the high conservation of the b-h subunit, questions remain about the evolutionary benefits of this fusion. Previous studies have suggested that subunit h may be involved the stability of the F_1_ sector along with the OSCP subunit and the C-terminus of the b-subunit (48). We were unable to identify subunit h in several lineages. While hmmsearches corroborated orthology between ASA4 and subunit h in green algae, no subunit h orthologues could be identified in red algae or glaucophytes. Similarly, while all other F_o_ subunits were identified in the Cryo-EM structures of *Trypanosoma brucei* and *Euglena gracilis*, subunit-h remains could not be identified. We modelled the b, d, h, 8, and OSCP subunits of several eukaryotes. Interestingly, the peripheral stalk proteins from Andalucia godoyi, Arabidopsis thaliana, or Phytophthora infestans were predicted to form realistic Type I structures using ColabFold (49) (Figure 6A). However, some structural aberrations in OSCP folding could be seen when the h subunit was omitted from the predictions (Figure 6B). Similarly, in lineages where the h subunit could not be found, the peripheral stalk did not form realistic Type I structures when modelled with ColabFold (Figure 6C). Given these structural predictions, subunit h appears to play an important structural role in Type I ATP synthases. Thus, it is possible that taxa in which we were unable to find an orthologue of subunit h may possess extremely divergent h-subunits or novel accretions that perform a similar function. By fusing subunit h to the C-terminus of subunit b in the SAR supergroup, the F_1_ sector may be more stable and thus be more efficient in ATP synthesis whilst losing its independent effectiveness as an ATPase – or, perhaps it was simply a stroke of luck.

**Figure 6.**
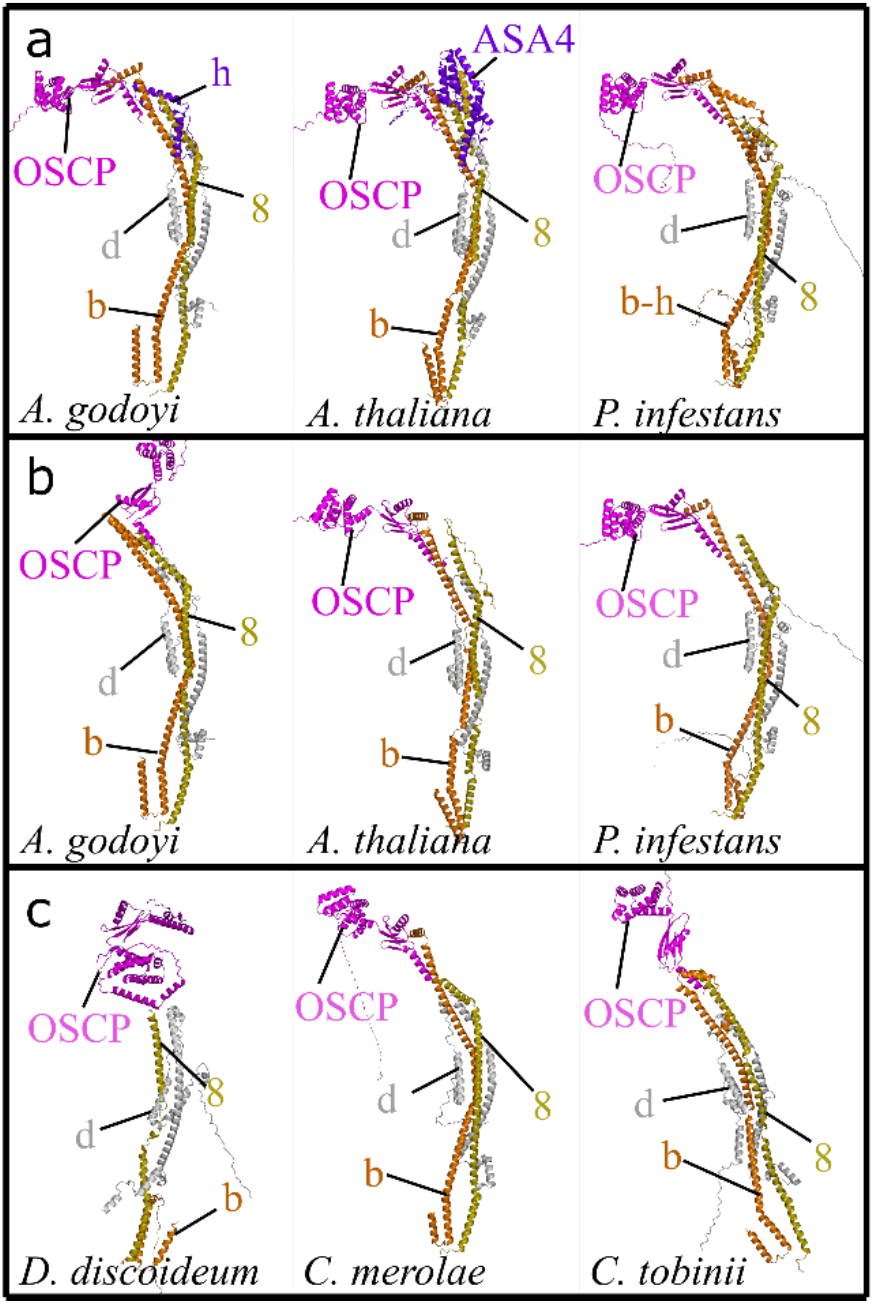
Multi-subunit modelling of ATP synthase peripheral stalk structures provides insight into subunit h function. Peripheral stalk structures were modelled from diverse eukaryotes **(a)**. *Andalucia godoyi*, *Arabidopsis thaliana*, and *Phytophthora infestans* subunits collected fold into believable peripheral stalk structures. **(b)**. Peripheral stalks from *Andalucia godoyi*, *Arabidopsis thaliana*, and *Phytophthora infestans* have difficulty folding correctly without subunit h. **(c)**. Predicted structures from *Dictyostelium discoideum*, *Cyanidioschyzon merolae*, and *Chrysochromulina tobinii* fold incorrectly, similar to structures lacking subunit h. Sequences from each organism were obtained from BLAST and HMMER into the EukProt3 and NCBI databases to obtain nuclear and mitochondrial genomes respectively (21–23, 25, 62). Peripheral stalk structures were assembled from our findings via the “ColabFold: AlphaFold2 using MMseqs2” notebook (28, 49). Structures with Type I-like b, d, h, and 8 subunits folded into structures resembling animal or fungal Type I ATP synthase (e.g., *Andalucia godoyi*, *Arabidopsis thaliana*, and *Phytophthora infestans*). Conversely, any structure with a divergent or missing h subunit could not be modelled into a believable peripheral stalk (e.g., *Dictyostelium discoideum*, *Cyanidioschyzon merolae*, *Chrysochromulina tobinii*).

### Conclusions

While the amino acid composition of ATP synthase subunits varies across eukaryotes, conservation of core fungal and animal subunits implies that the ancestral structure of LECA ATP synthase resembled the Type I complex present in animals and fungi. This would appear true for nearly any proposed rooting hypothesis (23, 50–56). The ancestral-like qualities of *Andalucia godoyi* ATP synthase and the apparent lack of dimerization subunits e and g in jakobids (Figure 2) is consistent with a Jakobid root (50), implying a subsequent addition of these subunits after the divergence of jakobids from all other eukaryotes. However, very few jakobids have associated sequence data and many more jakobids must be sequenced to strengthen this suggestion. The core subunit composition of ATP synthase is largely conserved in Type II, III, and IV ATP synthases as we and others identified subunits originally thought to be novel and lineage specific as divergent orthologues of ancestral subunits (Figure 2, Datasets S1-S4, and (14–18, 35, 57)). This indicates that accretion of novel subunits is more prevalent than replacement of ancestral subunits. In fact, very few replacements were detected, with the only notable instances including subunits e and g, which are completely replaced in some Type II structures and subunit h, which is possibly replaced in Type IV structures. A wide range of taxa that lack identifiable ancestral subunits are prime targets for future structural studies. These include amoebozoans (lack subunits h and k), jakobids/tsukubamonads (lack subunits e and g), heteroloboseans (lack subunits b, f, h, i/j, and k), stramenopiles (lack subunits e and g), retarians (lack subunits b-h, f, 8, i/j, and k), other rhizarians (lack subunit f), haptophytes (lack subunits e, g, and h), and rhodophytes (lack subunit h).

The widespread conservation of subunits implies that virtually all uniquely mitochondrial features of ATP synthase (e.g., dimerization) evolved prior to the divergence of eukaryotes. Previous work showed that the Mitochondrial Contact Site and Cristae Organizing System (MICOS) originated from homologs in bacteria (58), and that mitochondrial cristae are likely homologous to alphaprotebacterial intracytoplasmic membranes (ICMs) (59). While cristae formation requires ATP synthase – and likely ATP synthase dimerization – it is unclear what is required for alphaproteobacterial ICM formation (60). Closer investigation of ATP synthases in diverse alphaproteobacteria will shed light onto early evolution of mitochondrial cristae.

## Materials and Methods

### Database selection and homology searching

Sequences from organisms with well-annotated genomes or solved structures of ATP synthases (e.g., *Saccharomyces cerevisiae*, *Homo sapiens*, *Arabidopsis thaliana*, *Acanthamoeba castellani*, *Polytomella parva*, *Tetrahymena thermophila*, *Toxoplasma gondii*, *Trypanosoma brucei*, and *Euglena gracilis*) were retrieved from the NCBI sequence database to be used in initial queries against a database of 219 EukProt3 (23) proteomes selected based on their taxonomic breadth and estimated genome completion (61). BLAST (21, 22, 62) (v2.12.0, word size 3) queries were conducted into 219 selected proteomes. Top BLAST hits with e values below a threshold of 0.01 were then used as reciprocal BLAST queries against the proteomes with experimentally validated orthologues. A protein sequence was considered a validated orthologue if the experimentally validated ATP synthase subunits were returned as top hits in the reciprocal BLAST searches. To detect divergent homologues, validated orthologues were aligned using MUSCLE (63) to make HMMs to search all predicted proteomes lacking validated orthologues using hmmsearch v3.3.2 (24, 25). Top hits from HMM searches were validated via reciprocal BLAST, phmmer, and HHpred v3.3.0 (64) searches into predicted proteomes with known ATP synthase orthologues. In some cases, putative orthologues could not be validated and an in silico structural approach was taken. Homology modelling tools like Phyre2 (26) and SWISS-MODEL (27) were initially used to identify proteins that have predicted structural similarities to ATP synthase components of S. cerevisiae and H. sapiens. Finally, AlphaFold2 (28) was used to predict the structures of divergent ATP synthase subunits whose orthology could not be validated by other means. These predicted structures were aligned with solved structures PyMOL v2.5 (65–67), to identify potential similarities in protein structure. All ATP synthase subunit EukProt accessions and sequences are in Datasets S1 and S3, respectively.

## Acknowledgments

This work was supported by the National Science Foundation, DBI-2119963, BII: Mechanisms of Cellular Evolution.

