## Supplemental Figures 1-9 for "The persistent homology of mitochondrial ATP synthases"

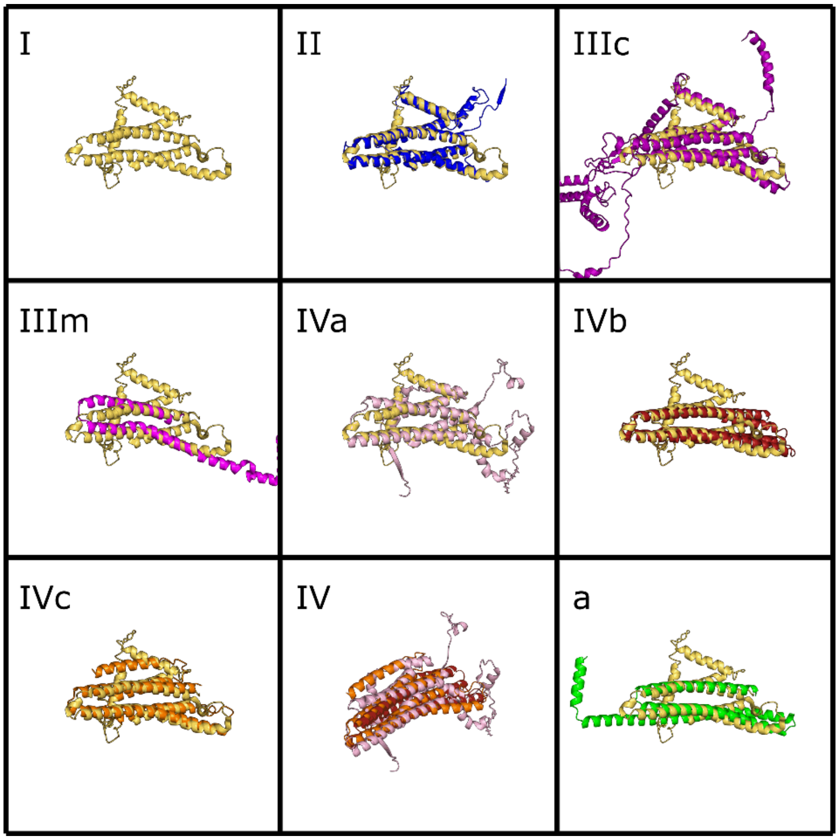


**Fig. S1. Variation in structure of subunit a across eukaryotes.** The C-terminus of the a-subunit which interacts with the c-ring is well-conserved across all known aerobic eukaryotes and is usually encoded in the mitochondrial genome. PDB structures were compared to AlphaFold2 predictions (1) and aligned via PyMOL2 (2–4). [I]Structure of yeast subunit a (PDB: 6B8H); [II]*Polytomella parva* subunit a (PDB: 6RD4); [IIIc]*Tetrahymena thermophila* subunit a (PDB: 6YNY); [IIIm]*Toxoplasma gondii* subunit a (PDB: 6TMK); [IVa]*Euglena gracilis* subunit a (PDB: 6TDU). [IVb]*Trypanosoma brucei* and [IVc]*Sulcionema specki* subunit a sequences were obtained from predicted *atp6* mitochondrial genes or Cryo-EM studies (5). [IV]Euglenozoan a-subunits overlayed. [a]*Globobulimina sp.* subunit a. The putative *Globobulimina sp. atp6* gene was identified using tBLASTn searches into the TSA database on NCBI (6–8). Yeast subunit in yellow.

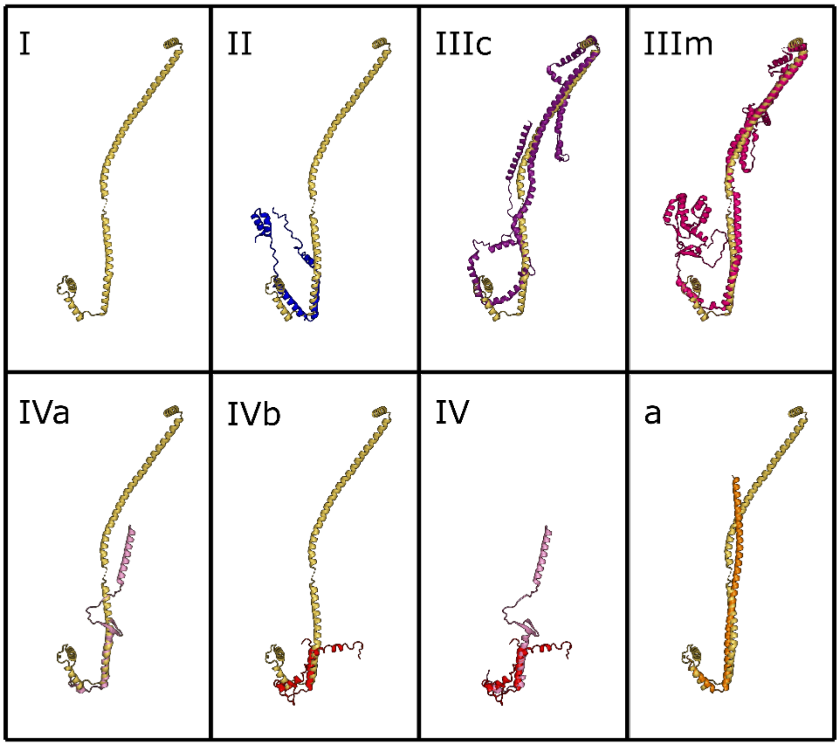


**Fig. S2. Variation in structure of subunit b across eukaryotes.** The structure of the N-terminus of the b-subunit is conserved across divergent eukaryotes. Solved structures were obtained from PDB: [I]Structure of yeast subunit b (PDB: 6B8H); [II]*Polytomella parva* subunit ASA6 (PDB: 6RD4); [IIIc]*Tetrahymena thermophila* subunit b (PDB: 6YNY), [IIIm]*Toxoplasma gondii* subunit b (PDB: 6TMK); [IVa]*Euglena gracilis* subunit b (PDB: 6TDU). Sequences from [IVb]*Trypanosoma brucei* and [a]*Acanthamoeba castellani* were taken from published reports (5, 9). These protein sequences were structurally modelled using AlphaFold2 (1). [IV]Euglenozoan b-subunits overlayed. Structures were aligned using PyMOL2 (2–4). Yeast subunit in yellow.

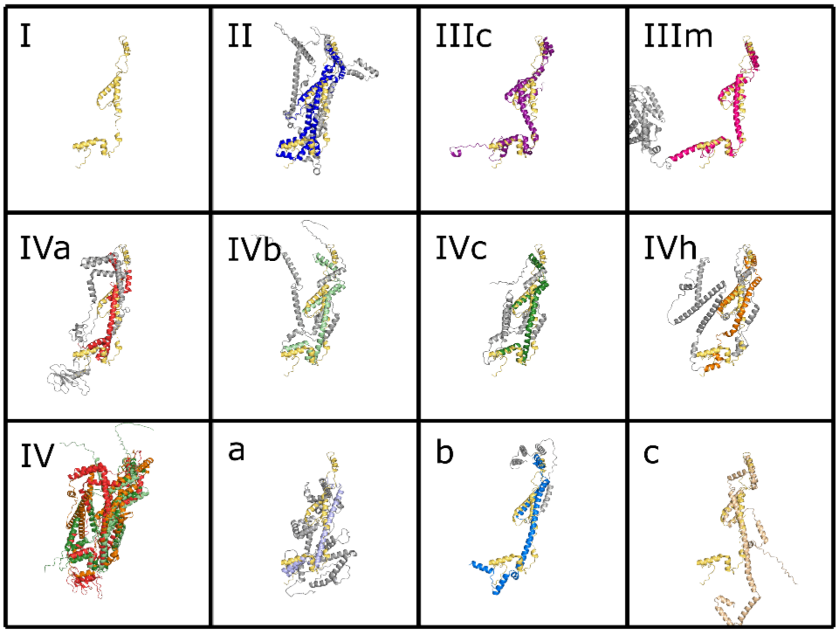


**Fig. S3. Variation in structure of subunit d across eukaryotes.** The C-terminus helices are well conserved even across divergent orthologues of the d-subunit. Solved structures were obtained from PDB: [I]Structure of yeast subunit d (PDB: 6B8H); [II]*Polytomella parva* (PDB: 6RD4); [IIIc]*Tetrahymena thermophila* (PDB: 6YNY); [IIIm]*Toxoplasma gondii* (PDB: 6TMK); and [IVa]*Euglena gracilis* (PDB: 6TDU). Sequences from [IVb]*Trypanosoma brucei* and [a]*Acanthamoeba castellani* were taken from published reports (5, 9–11). Orthologues identified in [IVc]*Rhynchopus humris*, [IVh]*Naegleria gruberi*, [b]*Thecamonas trahens*, and [c]*Reticulomyxa filosa* were acquired via BLAST and hmmsearch (6, 8, 12). [IV]Discicristate d subunits overlayed. The structures of these unmodelled proteins were predicted using AlphaFold2 (1) and aligned with other orthologues through PyMOL2 (2–4). Yeast subunit in yellow. Novel N and C-terminus extensions are shown in gray.

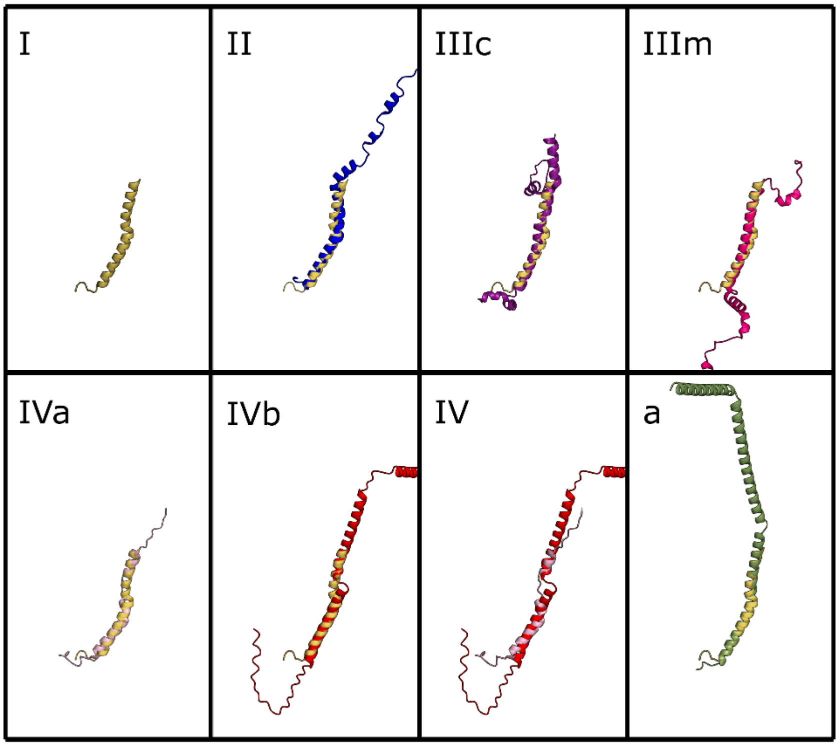


**Fig. S4. Variation in structure of subunit 8 across eukaryotes.** Subunit 8 is the orthologue of the bacterial subunit b' (13). Subunit 8 is truncated in animals and fungi and many other eukaryotic lineages but remains much longer in other eukaryotes. In *Andalucia godoyi*, subunit 8 is much longer, reminiscent of its bacterial counterpart subunit b'. Structural variations occur across all eukaryotes with unclear functional implications. Solved structures were obtained from PDB: [I]Structure from yeast (PDB: 6B8H); [II]*Polytomella parva* (PDB: 6RD4); [IIIc]*Tetrahymena thermophila* (PDB: 6YNY); [IIIm]*Toxoplasma gondii* (PDB: 6TMK); and [IVa]*Euglena gracilis* (PDB: 6TDU). The [IVb]*Trypanosoma* and [a]*Andalucia* proteins were acquired from their respective studies (5, 14) and modelled using AlphaFold2 (1). [IV]Euglenozoan 8-subunits were overlayed by alignment using PyMOL2 (2–4). Yeast subunit in yellow.

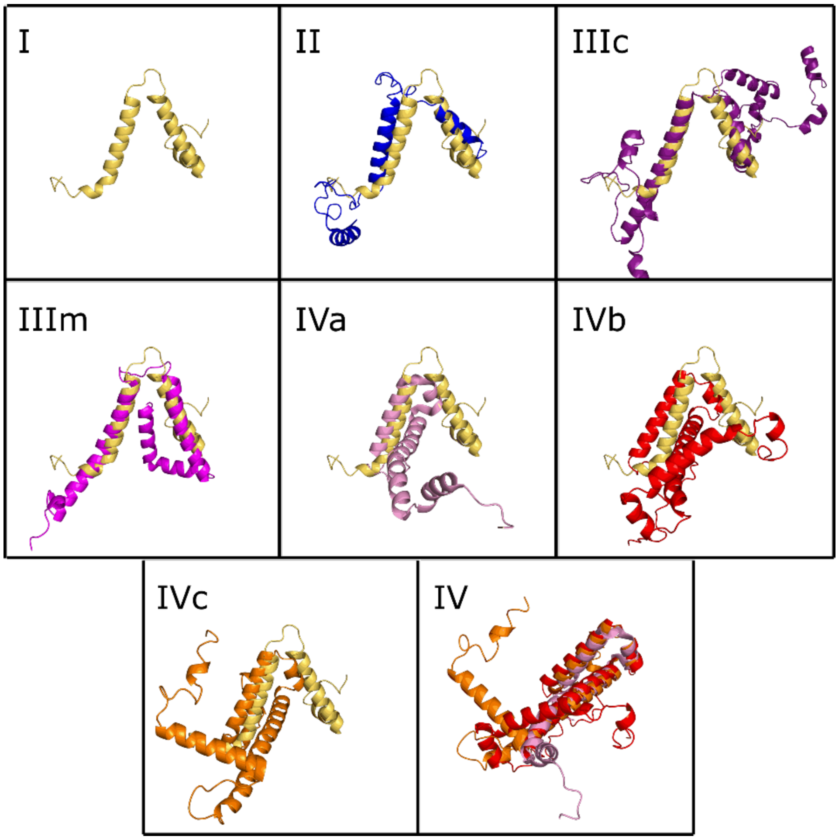


**Fig. S5. Variation in structure of subunit f across eukaryotes.** While the sequence of subunit f has diverged significantly among Types II-IV, its inverted V-shape structure remains well-conserved. Solved structures were obtained from PDB: [I]Structure from yeast (PDB: 6B8H); [II]*Polytomella parva* (PDB: 6RD4); [IIIc]*Tetrahymena thermophila* (PDB: 6YNY); [IIIm]*Toxoplasma gondii* (PDB: 6TMK); [IVa]*Euglena gracilis* (PDB: 6TDU). The [IVb]*Trypanosoma brucei* sequence was obtained from prior studies of kinetoplastid ATP synthase subunit composition (11) and was used to find orthologues in diplonemid [IVc]*Sulcionema specki* via HMMER (12). [IV]Euglenozoan f-subunits overlayed and rotated to show better alignment. Sequences were structurally modelled and aligned with yeast subunit f using AlphaFold2 and PyMOL2, respectively (1–4). Yeast subunit in yellow.

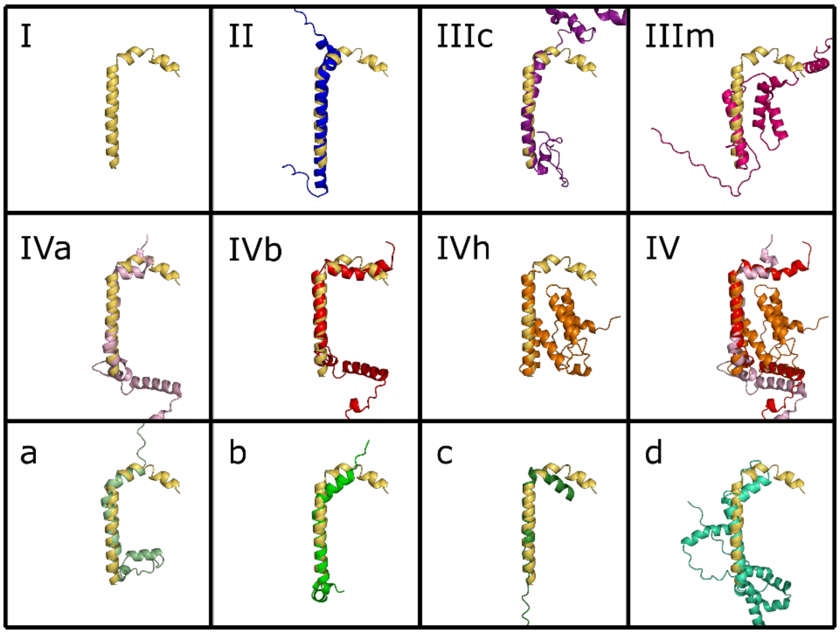


**Fig. S6. Variation in structure of subunit i/j across eukaryotes.** While the sequence of subunit i/j has diverged significantly among Types II-IV, its simple structure remains relatively well-conserved. Solved structures were obtained from PDB: [I]yeast (PDB: 6B8H), [II]*Polytomella parva* (PDB: 6RD4), [IIIc]*Tetrahymena thermophila* (PDB: 6YNY), [IIIm]*Toxoplasma gondii* (PDB: 6TMK), and [IVa]*Euglena gracilis* (PDB: 6TDU) were obtained from PDB. The [IVb]*Trypanosoma brucei*, [a]*Acanthamoeba castellani*, and [b]*Arabidopsis thaliana* were obtained from prior reports (5, 9, 11, 15). [IVh]*Naegleria gruberi*, [c]*Andalucia godoyi*, and [d]*Acanthocystis sp. HF-20* were identified through HMMER and HHpred (12, 16). [IV]Discicristate i/j subunits overlayed using PyMOL2 (2–4). All proteins from unsolved structures were modelled using AlphaFold2 (1). Yeast subunit in yellow.

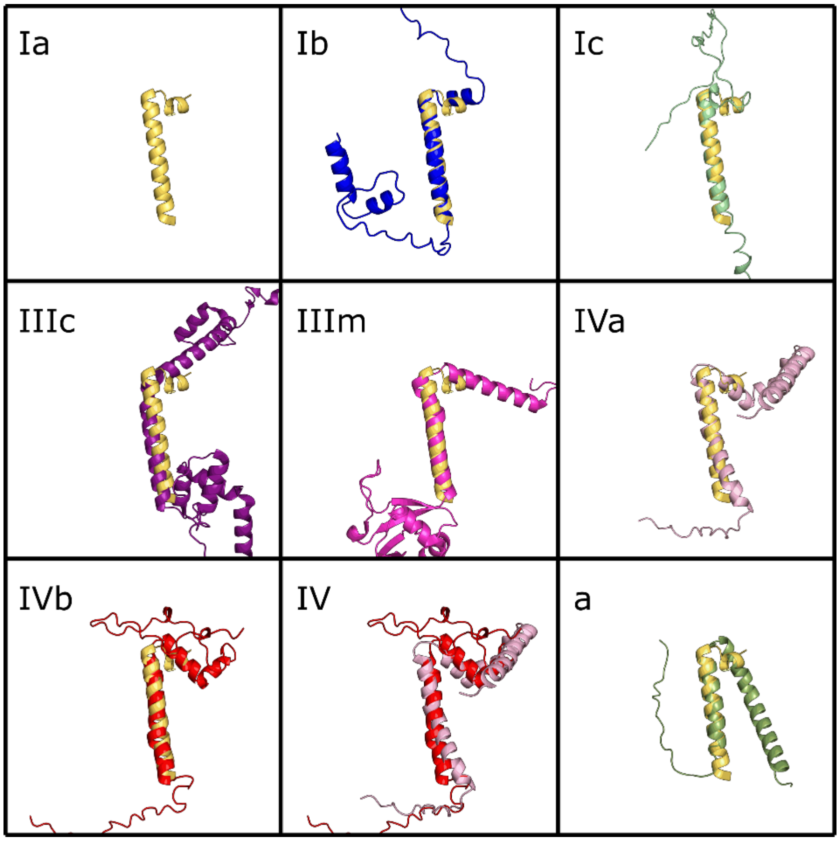


**Fig. S7. Variation in structure of subunit k across eukaryotes.** Similar to subunits f and i/j, the sequence of subunit k has diverged significantly among Types II-IV while its structure remains relatively well-conserved. Solved structures were obtained from PDB: [Ia]bovine (PDB: 6ZPO), [IIIc]*Tetrahymena thermophila* (PDB: 6YNY), [IIIm]*Toxoplasma gondii* (PDB: 6TMK), and [IVa]*Euglena gracilis* (PDB: 6TDU) were obtained from PDB. The [IVb]*Trypanosoma brucei* sequence was acquired from the Cryo-EM study of the kinetoplastid ATP synthase (5). Sequences from [Ib]*Ectocarpus siliculosus*, [Ic]*Glycine max*, and [a]*Andalucia godoyi* were found through BLAST and HMMER (6, 8, 12). All sequences from organisms without solved structures were modelled through AlphaFold2 and aligned via PyMOL2 (1–4). Bovine subunit in yellow.

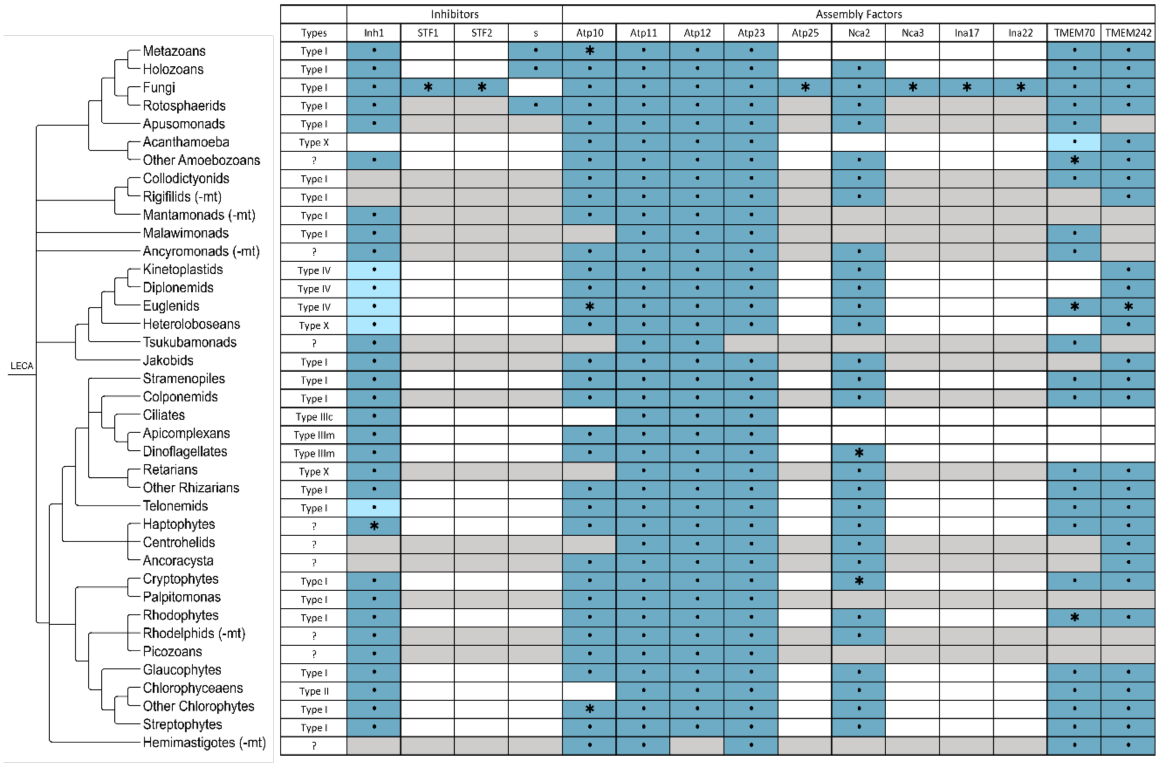


**Fig. S8. Most eukaryotes retain a core set of ancestral assembly factors.** Some putative assembly factor orthologues were identified in 219 predicted proteomes from diverse eukaryotic lineages (See Dataset S1 for accessions). Through BLAST and HMMER queries (6, 8, 12), we collected sequences from the EukProt3 database (17). Ancestral-like subunits were verified through reciprocal BLAST or phmmer are shown in dark blue (6, 8, 12). Divergent orthologues verified through HHpred or structural modelling are coloured light blue (1, 16, 18). •These subunits are encoded in the nucleus. *Subunits are only present in certain lineages.

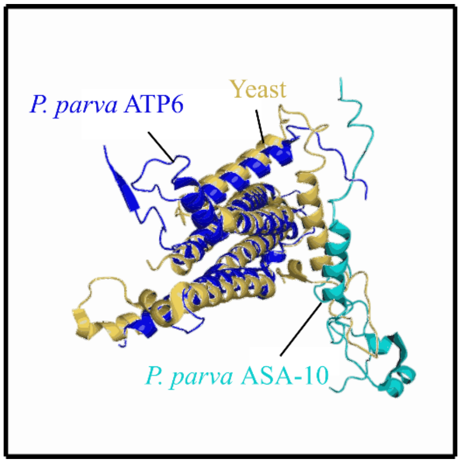


**Figure S9. ASA10 replaces the N-terminal helix of eukaryotic subunit a.** The *Polytomella* subunit a structure lacks an N-terminal helix conserved across most eukaryotes, being entirely replaced by the ASA10 subunit. Structures for the yeast subunit a (PDB: 6B8H) and the *Polytomella* subunit a – ASA10 complex (PDB: 6RD4) were obtained from PDB and aligned using PyMOL2 (2–4). Yeast subunit a is represented by the yellow structure and *Polytomella* subunits a and ASA10 are depicted by the dark and light blue structures, respectively.

**Dataset S1 (separate file). ATP synthase subunit and assembly factor accessions from 219 EukProt3 Genomes and Transcriptomes.** All ATP synthase subunits are well-conserved across most lineages with some notable absences among subunits e, f, g, h, and k. These sequences were acquired via extensive BLAST and HMMER searches into each genome and transcriptome in which top hits validated through reciprocal BLAST, phmmer, HHpred, or structural modelling techniques were recorded (1, 6, 8, 12, 16, 18, 19). Sequences from mitochondrial genomes or transcriptomes absent in EukProt3 were acquired from NCBI via BLASTp or tBLASTn respectively (6–8).

**Dataset S2 (separate file). Most Type II-IV ATP synthase subunits are widely conserved within their select lineages.** To determine the conservation of novel subunits across taxa, BLAST and phmmer searches using reference sequences from previously studied organisms (*Polytomella parva*, *Tetrahymena thermophila*, *Toxoplasma gondii*, *Trypanosoma brucei*, and *Euglena gracilis* (5, 20–23)) were utilized to identify orthologues across other members of their respective taxa (6, 12). Top hits with E-value < 0.01 that could be verified via reciprocal BLAST and phmmer were recorded (6, 8, 12).

**Dataset S3** (https://doi.org/10.6084/m9.figshare.21027730)**. ATP synthase core subunit, inhibitor, and assembly factor FASTA files.** Sequences from solved structures (5, 20–23) were used to conduct BLAST (6, 8) and HMMER (12) searches into 219 organisms from the EukProt v3 database (17) and tBLASTn (7) queries into the NCBI TSA, WGS, and SRA databases. These hits were verified via reciprocal BLAST (6, 8), phmmer (12), HHpred (16), Phyre2 (19), SWISS-MODEL (18), and AlphaFold (1).

**Dataset S4** (https://doi.org/10.6084/m9.figshare.21056146)**. Novel ATP synthase subunit FASTA files for Types II-IV.** Sequences from novel structures (5, 20–23) were used to conduct BLAST (6–8) and HMMER (12) searches into closely related genomes and transcriptomes obtained from the EukProt v3 database (17). These hits were verified via reciprocal BLAST (6, 8), phmmer (12), HHpred (16), Phyre2 (19), SWISS-MODEL (18), and AlphaFold (1).
